# Cancer stem cell transcriptomic signatures reveal patient-to-patient heterogeneity emphasizing the need for patient-specific ovarian cancer targets

**DOI:** 10.64898/2026.09.30.755773

**Authors:** Matthew S. Jung, Lindsey J. Davis, Audrey A. Birdwell, Jisun Kang, Colleen Feng, Amelia Pi, Sue Chin Lee, Sandra Davern, Rachel Patton McCord, Junming Yue, Wenjing Zhang, Won-Young Choi, Rachel S. Perkins

## Abstract

**Background/Objectives:** In high-grade serous ovarian cancer (HGSOC), the abilities of tumor cells to resist cytotoxic treatment and relapse are largely driven by ovarian cancer stem cells (CSCs). CSCs are a population of cells within a tumor that maintain stem-like qualities and have the capacity to produce metastasis and drive drug resistance. Based on the complexity of CSC signaling and emerging evidence of differential isoform usage in development, we hypothesize that CSCs possess a patient-specific, heterogeneous landscape which exploits isoform diversity to sustain stemness, metastasis and drug resistance.

**Methods:** RNA-seq was generated from 3 patient-derived HGSOC (PD-HGSOC) to investigate stemness and differentiation status comparing standard culture and CSCs. The data was analyzed at both gene- and isoform-levels to identify patient-specific CSC features and define mechanisms of cancer-related stemness and plasticity.

**Results:** CSCs exhibited higher stemness characteristics and identified differentially expressed gene sets across PD-HGSOCs. We noted that each patient had a distinct subset of genes regulating stemness phenotypes which suggested multiple mechanisms of CSC development and maintenance. Additionally, a large proportion of gene-level expression changes were driven by the differential expression of a single predominant isoform. Importantly, our isoform-level analysis enabled the identification of differential isoform usage that would not be detected with gene level expression analysis.

**Conclusions:** CSC phenotypes such as chemoresistance, metastasis and self-renewal are conserved across patients, but arise from largely patient-specific gene and isoform programs. This CSC heterogeneity is reflective of the heterogeneity seen in patients and should be further evaluated in deriving precision-medicine approaches for HGSOC.

**Graphical Abstract:** 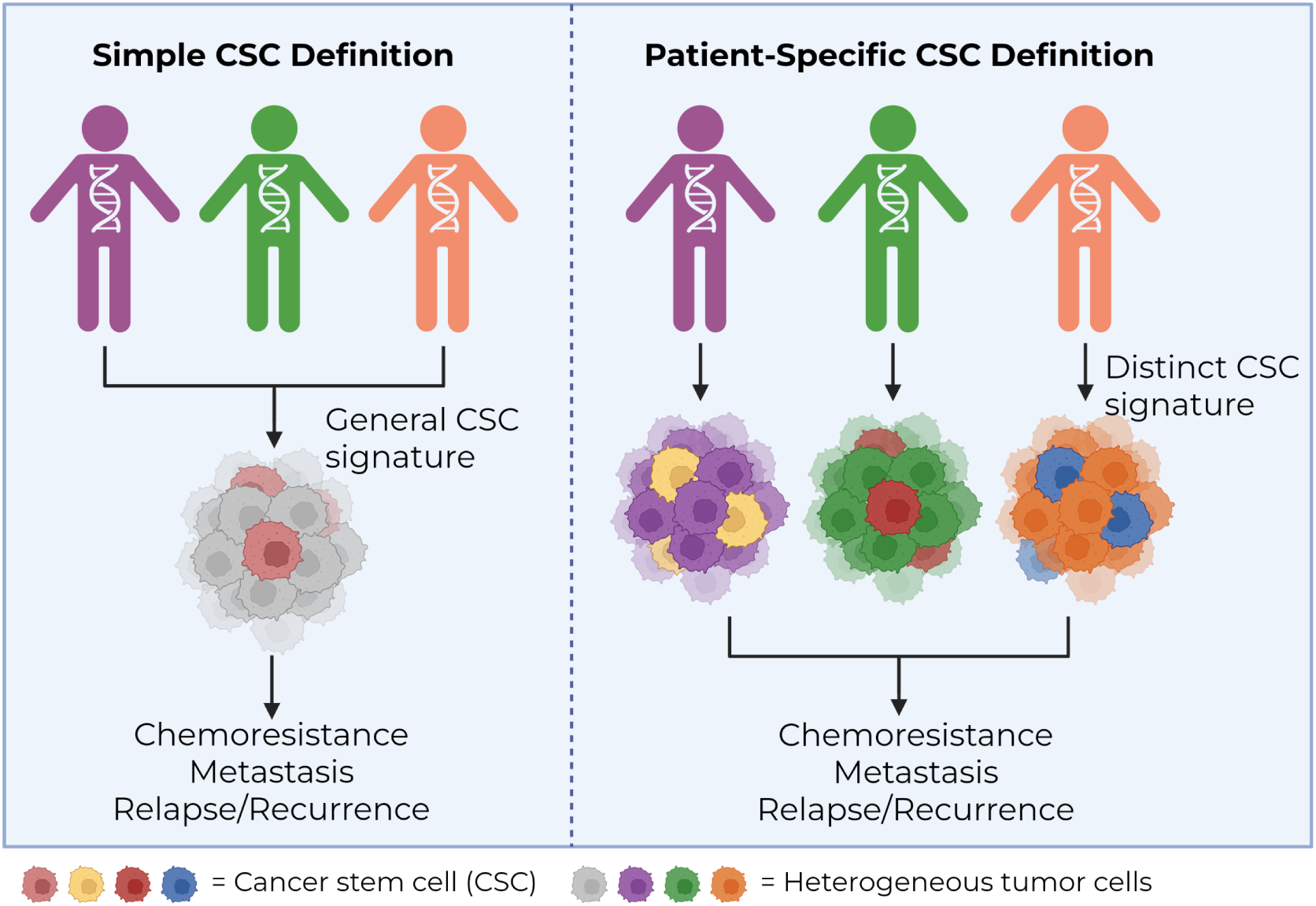

Simple vs Patient-Specific CSC signatures. Created in BioRender. Perkins, R. (2026) https://BioRender.com/m8qwrmt

**Simple Summary:** Patients with advanced ovarian cancer have poor prognosis, no targeted treatment options and are often diagnosed at late stages due to rapid metastasis and vague symptoms. Cancer stem cells contribute to these poor patient outcomes via increased drug resistance, tumor relapse, and metastasis. Here we found that cancer stem cell phenotypes such as chemoresistance, metastasis and self-renewal are conserved across patients, but arise from largely patient-specific gene and isoform programs. This heterogeneity is reflective of the heterogeneity seen between patients and should be further evaluated in order to develop precision-medicine approaches for ovarian cancer. Our results provide foundational understanding of cancer stem cell plasticity, patient heterogeneity, and next generation drug design, increasing target specificity, and reducing adverse effects.

## 1. Introduction

Ovarian cancer remains the most lethal gynecologic malignancy, largely because most patients present with advanced disease and, despite an initial response to cytoreductive surgery and platinum-based chemotherapy, eventually develop recurrent and treatment-resistant tumors [1–4]. High-grade serous ovarian cancer (HGSOC), the most common and aggressive histologic subtype, is characterized by extensive intratumoral and interpatient heterogeneity, widespread peritoneal dissemination, and a remarkable capacity to adapt to therapeutic pressure [5,6]. These features present a major challenge for the development of durable targeted therapies because tumors with similar histopathologic diagnoses may depend on substantially different molecular programs for progression, metastasis, and treatment resistance.

Cancer stem cells (CSCs) are a distinct subpopulation of cells within a tumor that possess stem-like properties, including self-renewal, pluripotency, and enhanced survival mechanisms [7]. These properties are thought to arise, at least in part, through aberrant reactivation of oncofetal and embryonic signaling programs that are normally restricted to early development [8]. A growing body of evidence indicates that CSCs are central drivers of the clinical challenges that limit the efficacy of conventional surgical, radiologic, and chemotherapeutic interventions [7]. These cells promote therapeutic resistance through elevated drug efflux and detoxification activity—mediated by ATP-binding cassette transporters and aldehyde dehydrogenases—as well as through slow cycling kinetics and the capacity to enter a quiescent state that renders them refractory to cytotoxic agents dependent on active cell division. CSCs further facilitate metastatic dissemination via induction of the epithelial-to-mesenchymal transition (EMT), a process increasingly understood to be mechanistically intertwined with CSC state acquisition and governed by overlapping regulatory circuits [9]. Following clinical intervention, surviving CSCs can reinitiate tumor growth through asymmetric division, underlying disease recurrence and relapse. This behavior is shaped by a highly interconnected regulatory network: the tumor microenvironment (TME). The TME establishes dynamic stem cell niches through bidirectional signaling, metabolic plasticity and convergent developmental and oncogenic signaling [8]. Together, these properties establish CSCs as both a major barrier to durable clinical response and a compelling therapeutic target. Consequently, resolving the coordinated mechanisms governing CSC regulation and identifying reliable CSC-specific markers remain essential prerequisites for the development of effective, CSC-directed therapeutic strategies.

A critical gap in knowledge in the pursuit of treatments directed towards CSCs, is the lack of a well-established signature spanning across cancers. In fact, it is well known that CSC markers vary between cancers [10]. It is becoming quite evident that classic CSC markers are insufficient to fully understand the heterogeneous landscape of cancer stemness. Furthermore, there is a lack of transcriptomic studies specifically characterizing CSCs isolated from patient-derived cells. Instead, many studies focus on constructing stem-cell signatures from publicly available patient whole-tumor sequencing, which does not capture the depth required to fully characterize this tumor driving population of cells. This study isolates and cultures CSCs from three HGSOC patient-derived cells to evaluate patient-specific CSC transcriptomic landscape. We show that CSCs from patient to patient have a unique transcriptional signature with conserved phenotypic outcomes including upregulation of chemoresistance pathways, metastatic pathways and oncofetal/embryonic pathways. Additionally, the limited number of conserved genes between all three patients does not reflect an upregulation of the classic stemness markers, instead they highlight the complex and dynamic involvement of CSCs in the regulation of the extracellular matrix, growth factor signaling and interferon response pathways.

## 2. Materials and Methods

### Patient derived ovarian cancer culturing and cancer stem cell enrichment

Patient-derived ovarian cancer cells (PDCs) were purchased from the NCI’s Patient Derived Material Repository (PDMR) under material transfer agreement. The three PDCs used for this manuscript are from patients with HGSOC. 1. HG = Primary HGSOC from the right ovary of a 74-year-old treatment naïve patient (575813-281-R2-J2-PDC). 2. PM = Peritoneal metastatic HGSOC from a peritoneal biopsy of an 81-year-old treatment naïve patient (994656-043-R-J2-PDC). 3. UM = Peritoneal metastatic HGSOC from an anterior umbilical biopsy of a 63-year-old multidrug resistant patient (655913-031-T-J1-PDC) (**Figure 1A-C**). HG, PM, and UM were grown in accordance with the standard operating procedures provided by the NCI’s PDMR (SOP30101, SOP30102, SOP30103, SOP30104). Briefly, PDCs were cultured in complete media containing Advanced DMEM/F-12 (Gibco^TM^, Thermo Fisher Scientific, U.S.), heat-inactivated fetal bovine serum (Cytiva HyClone^TM^, U.S.), recombinant Human EGF (R&D Systems^TM^, U.S.), Hydrocortisone (Sigma Aldrich, U.S.), Adenine (Sigma Aldrich, U.S.), Y-27632 dihydrochloride (MedChemExpress, U.S.), GlutaMAX (Gibco^TM^, Thermo Fisher Scientific, U.S.) and Primocin (InvivoGen, U.S.). According to NCI recommendations, HG and PM PDCs were maintained in LDEV-free Matrigel^TM^ (Corning^TM^, U.S.) coated flasks, and UM PDCs were maintained in non-coated flasks at 37C° and 5% CO_2_. The culturing method described above maintained cells in adherent (ADH) or “bulk” population state. To enrich for cancer stem cells (CSCs), PDCs were removed from attachment, plated in non-adherent tissue culture plates and their media was changed to serum-free DMEM (Gibco^TM^, Thermo Fisher Scientific, U.S) and supplemented with B27 (Gibco^TM^, Thermo Fisher Scientific, U.S), recombinant Human bFGF (R&D Systems^TM^, U.S.), recombinant Human EGF, and Primocin. This culturing condition formed 3D-CSC-enriched spheroids (SPH) which is a well-established model of CSCs [11,12].

**Figure 1.**
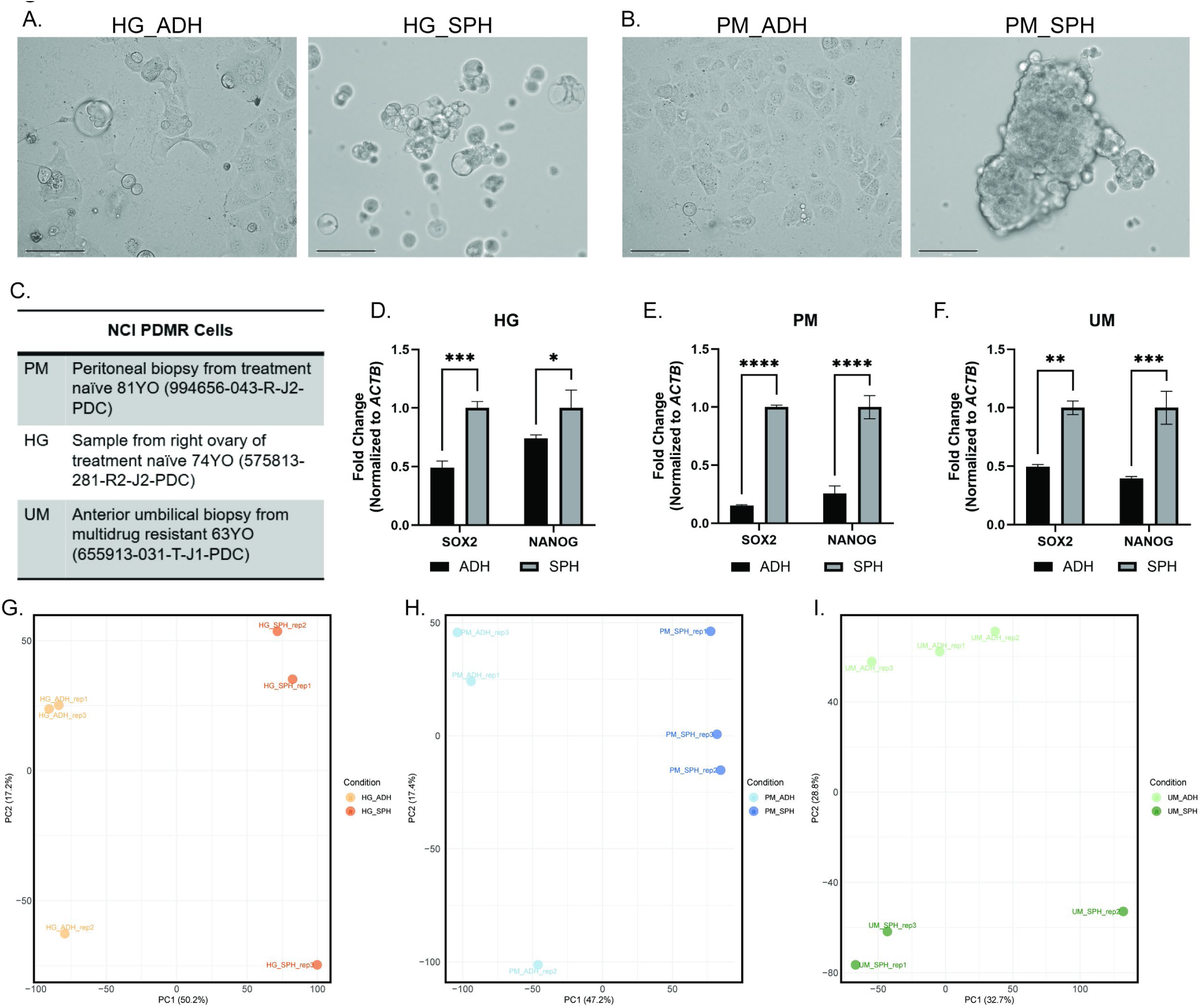
Establishment and characterization of cancer stem cell (CSC) models. A-B) Representative 20X images of HG (A) and PM (B) ADH vs SPH image, scalebar = 125 µm. C) Table of patient data for the patient derived cells used in this study. Cells from NCI PDMR. D-F) RT-qPCR analysis of stemness markers *SOX2* and *NANOG* in ADH vs SPH cultures across HG, PM, and UM. (G-I) PCA of RNA-seq data from PDCspecific PCA plots showing transcriptomic separation between ADH and SPH conditions in HG (G), PM (H), and UM (I) cells individually.

### RNA-seq sample generation and purification

PDCs were plated in either ADH or SPH conditions as described above. Prior to collection for RNA-seq, RT-qPCR was performed to ensure the appropriate markers (*SOX2*, *NANOG*) were increasing within CSCs compared to their ADH counterparts. Then, HG, PM, and UM PDCs were plated in triplicate wells of a 6-well adherent and non-adherent plate to enrich for whole ADH population or CSC population. After 72 hours, ADH and SPHs were collected and RNA was isolated using TRIzol^TM^ Reagent (according to manufacturer’s recommendations; Invitrogen^TM^, U.S.). RNA quantification was determined with NanoDrop and the quality was assessed using TapeStation to verify that each sample’s RIN value was >6. Samples were then sent to Azenta for sequencing.

### RNA-seq data generation and processing

RNA-seq data were generated by Azenta using their Standard RNA-Seq service with poly(A) selection and Illumina paired-end sequencing. Paired-end reads were aligned to the human reference genome (GRCh38/hg38) using STAR (version 2.7.11b). The aligner was configured to allow up to 20 multiple alignments per read (--outFilterMultimapNmax 20) and to require a minimum splice junction overhang of 8 nucleotides (--alignSJoverhangMin 8). The maximum number of mismatches was constrained to 10% of the read length (--outFilterMismatchNoverLmax 0.1), and introns were restricted to a size range of 20 to 1,000,000 bp (--alignIntronMin 20, --alignIntronMax 1000000; --alignMatesGapMax 1000000). Read filtering was based on detected splice junctions (--outFilterType BySJout), with minimum alignment score and matched-length thresholds scaled to read length (--outFilterScoreMinOverLread 0.33, --outFilterMatchNminOverLread 0.33). STAR produced unsorted BAM alignment files (--outSAMtype BAM Unsorted), retained unmapped reads within the BAM output (--out-SAMunmapped Within), and generated gene-level read counts (--quantMode GeneCounts).

### Identification of differentially expressed genes

Differential expression was evaluated with the DESeq2 package (v 1.38.3). Gene-level count tables were first restricted to protein-coding entries, and features located on chromosome Y were removed. Genes with fewer than five reads summed across all samples were excluded, and entries with duplicated gene symbols were collapsed so that each protein-coding gene was represented only once. The resulting matrix was used to build a DESeq2 dataset with experimental condition (HG_ADH, HG_SPH, PM_ADH, PM_SPH, UM_ADH, UM_SPH) specified as the design factor. Pairwise contrasts between conditions were then computed, and genes were considered differentially expressed if they passed a Benjamini-Hochberg-adjusted P value threshold of 0.05. For downstream visualization, transcript-per-million (TPM) values were transformed as log2(TPM + 1), centered and scaled by gene (z-score), and subjected to hierarchical clustering with Ward’s linkage, as implemented in ComplexHeatmap (v. 2.14.0). Annotated heatmaps were produced to highlight expression patterns and condition-specific signatures. A subset of robust DEGs, defined by padj ≤ 0.05 and an absolute log2 fold change greater than 2, was carried forward for pathway-level interpretation. Pathway specific heatmaps were generated with the following gene sets: Chemoresistance [13–15], Metastasis [16,17], Oncofetal/Embryonic/Stemness [18–20] and GO:0048568. Functional enrichment and comparative co-enrichment of these gene sets were examined using ToppCluster (https://toppcluster.cchmc.org/), applying a hypergeometric test with Benjamini–Hochberg correction and retaining terms with a false discovery rate below 0.05.

### Protein-protein interaction network analysis

Protein–protein interaction networks were constructed in STRING v12.0 (https://string-db.org/) using the full network type and a minimum required interaction score of 0.400. Edge confidence is given by the combined score, which integrates the neighborhood, gene fusion, co-occurrence, co-expression, experimental, database and text-mining channels; the separately reported homology score is not included in the combined score and was retained only to flag interactions between sequence paralogs, for which evidence may have been transferred between family members. Networks were partitioned into functional modules using the Markov cluster (MCL) algorithm implemented in STRING, with an inflation parameter of 3, and the resulting modules were used to identify pathway-level structure within each gene set. Functional enrichment of network modules was computed using a hypergeometric test against the whole-genome background with Benjamini–Hochberg correction, and terms with a false discovery rate below 0.05 were reported.

### Isoform switching analysis

Isoform switching between SPH and ADH cultures was analyzed separately for each PDC line (PM, HG, UM) using IsoformSwitchAnalyzeR in R, based on StringTie transcript-level quantifications and GENCODE v48 (hg38) gene and transcript annotations. After importing isoform counts and abundances and constructing a SwitchAnalyzeRlist object, lowly expressed isoforms were removed (gene and isoform expression >1, isoform fraction >0.01), and single-isoform genes were excluded. Differential isoform usage between SPH and ADH was tested with the DEXSeq framework (isoform_switch_q_value value < 0.05 and |dIF| > 0.05), followed by ORF prediction, intron retention assessment, and consequence analysis focusing on intron retention, nonsense-mediated decay, and ORF similarity. High-confidence coding-to-coding switches were defined as protein-coding isoforms without premature termination codons or intron retention that showed reciprocal isoform fraction changes within the same gene, and were summarized using IsoformSwitchAnalyzeR’s consequence plots, bar plots of gained versus lost isoforms per PDC, and heatmaps of dIF values across lines.

### RNA-seq validation

RNA-seq results were validated using RT-qPCR. cDNA was constructed from HG, PM, and UM ADH and SPHs at a concentration of 2000 ng using Thermo Scientific^TM^ Maxima First Strand cDNA Synthesis Kit. Then cDNA was utilized for RT-qPCR using SYBR Green Master Mix (ThermoFischer Scientific). The RT-qPCRs were performed on a QuantStudio 6flex PCR machine (Applied Biosystems) with SYBR Green cycling conditions set at 95°C for 10 min, 40 cycles of 95°C for 15 s and 60°C for 1 min, followed by detection of the melt curve. Primer designs for the RT-qPCR assays are reported in **Supplementary Table 1**.

RT-qPCR analysis calculated fold change between samples for relative gene expression. The cycle threshold (CT) value is used to calculate the sample fold change with the formula 2^-ΔΔCT^. The average fold change value and standard deviation are normalized to the housekeeping gene β-Actin (*ACTB*) to ensure equal amounts of cDNA are compared. The results are visualized in GraphPad and One-Way Anova or T-test is used to calculate significance between groups.

### Functional assays comparing ADH and SPH

#### Sphere Formation Assay

ADH PDCs were cultured in 6-well, Matrigel^TM^ coated, adherent plates and SPH PDCs were cultured in 6-well non-adherent plates. After 72 hours, ADH and SPH cells were collected, enzymatically separated with 0.05% Trypsin (Gibco, U.S.) at 37C° for 10 minutes. Cells were quenched and single cell sorted through 35 µm filter caps (Stellar Scientific, U.S.). Single cell suspensions were plated in SPH conditions in decreasing densities and allowed to grow for 72 hours. Plated ADH cells are primary SPHs and plated SPHs are secondary SPHs. SPH number was then counted by hand and plotted using GraphPad Prism Software.

#### Drug Response

ADH and SPH PDCs were cultured in 96-well adherent and non-adherent plates, respectively. After 72 hours, cells were treated with an increasing dose range of paclitaxel (PTX). 72 hours later, cell viability assays were performed according to manufacturer protocol using CellTiter-Glo® 3D Cell Viability Assay (Promega, U.S.). After incubation, each well was transferred to an opaque white-walled 96-well plate to measure luminescence. Results were analyzed and plotted using GraphPad Prism Software.

#### Migration

SPH PDCs were cultured in 6-well non-adherent plates. After 72 hours, SPHs were collected and separated into single cell suspensions as described above. In adherent 24 well plates with 8 µm pore transwell inserts (Nunc^TM^, Thermo Fisher Scientific, U.S), 500µL of complete media were added to the lower chamber, and SPH cells were plated in 200 µL serum and growth factor free DMEM in the upper chamber. Cells were incubated at 37C° and 5% CO_2_ for 20 hours before staining. After the incubation period, inserts were fixed and stained for imaging and analysis. Inserts were fixed in 500 µL of chilled 4% PFA for 15 minutes. Inserts were then submerged in 500 µL of 0.1% crystal violet for 15 minutes, washed in 1x PBS and then the non-migrated cells on the apical side of the chamber were removed with a cotton ball. Following staining, the inserts were left to air-dry. Four representative images, one from each quadrant of the insert, were acquired at 4X magnification using an EVOS microscope. Images were analyzed in ImageJ to quantify the area of the insert covered by crystal violet-stained cells.

## 3. Results

### Patient-derived ovarian cancer cell lines generate a robust CSC-like state under spheroid culture conditions

Ovarian cancer is characterized by extensive peritoneal dissemination and frequent relapse following chemotherapy, outcomes that are at least partly driven by a subpopulation of cells with CSC-like properties [21,22]. To investigate how CSC biology is shaped by tumor origin, we employed three ovarian cancer patient-derived cell lines (OV-PDCs) representing clinically distinct disease states including a high-grade primary tumor (HG), a peritoneal metastasis (PM), and an umbilical metastasis (UM). Under spheroid-forming conditions, OV-PDCs efficiently formed non-adherent spheroids, illustrating a clear morphological transition that has been widely used as a reliable in vitro readout for CSC enrichment in ovarian cancer (**Figure 1A-B**). Notably, the UM spheres are cultured in a mixed adherent/non-adherent culturing system as per NCI PDMR recommendations. Therefore, the selection for this OV-PDC is largely dependent on the transition to ultra-low attachment plates and CSC media, and therefore are morphologically similar between bulk cell cultures and CSCs. The three OV-PDCs consist of a wide range of patient characteristics including primary, metastatic, chemoresistant and treatment naïve (**Figure 1C**). This patient population, while limited by sample number, provides relevance as each OV-PDC captures the various clinical presentations of HGSOC (**Figure 1C**). Upregulation of established stemness markers (*SOX2*, *NANOG*) upon CSC induction was confirmed by RT-qPCR across all three lines indicating that our culture system reproducibly enriches for a CSC-like transcriptional state at the molecular level (**Figure 1D-F**). Principal component analysis of bulk RNA-seq profiles showed that adherent (ADH) and cancer stem cell (SPH) samples segregated along the primary axes of variation within each OV-PDC indicating that CSC formation is accompanied by a large-scale reorganization of the transcriptome rather than a modest quantitative shift in a limited number of genes (**Figure 1G-I**). Furthermore, a three-dimensional PCA including all 18 samples revealed that the three OV-PDCs occupy distinct transcriptional spaces even before CSC induction with ADH from the same PDC forming neighboring yet clearly separated clusters (**Supplementary Figure 1A**). Despite intrinsic differences between the different OV-PDC sources captured by PC1 and PC2, PC3 captures gene expression differences that consistently separate ADH and SPH states across all samples **(Supplementary information 1)**. This clustering pattern further enforces that the CSC culturing methods applied do not artificially enrich for one cell type (as would be seen if all three sets of CSCs clustered together), instead, it pulls the CSC subpopulation from its respective ADH PD-OVC culture – as seen by each ADH and SPH PD-OVC clustering individually and relatively close to each other. Taken together, these data show that transcriptional diversity reflects both cell-of-origin identity (HG, PM or UM) and cell subtype (ADH or SPH), providing a framework that allows us to investigate whether CSC identity is universal or molded by the tumor context from which it arises.

### OV-PDCs retain patient-specific transcriptional programs reflecting their clinical origins

As a prerequisite for comparing CSC states, we characterized the three OV-PDCs in their baseline adherent condition to confirm that each line retains hallmarks consistent with its clinical origin. Metastasis-associated genes including *ANGPTL4*, *MMP2* and *LGR5* were significantly upregulated in both PM and UM relative to HG, while *MMP1* was induced in PM only (**Figure 2A-D**), consistent with their reported roles in ovarian cancer metastasis. *ANGPTL4* promotes ovarian cancer cell migration, invasion and early peritoneal colonization in vivo [23]. *MMP1* and *MMP2* facilitate mesothelial barrier disruption and peritoneal dissemination by degrading extracellular matrix and injuring mesothelial cells [24,25]. *LGR5* functions as a stem/progenitor marker in the ovary and is overexpressed in epithelial ovarian cancer where it enhances proliferation, invasion and metastatic potential [26,27]. Chemoresistance-associated markers *NTSR1* and *SNCG* were most highly expressed in the UM PDC, consistent with reported roles for neurotensin and synuclein-γ signaling in acquired platinum resistance in ovarian cancer (**Figure 2E-F**). These RNA-seq observations were validated by RT-qPCR confirming that the clinical identity of each PDC is preserved in vitro (**Supplementary Figure 1B**). To characterize the transcriptional differences existing among the three OV-PDC lines, we performed pairwise differential expression analysis across HG, PM and UM PDCs grown under ADH culture condition and applied unsupervised k-means clustering to the resulting gene sets. This analysis revealed distinct gene clusters selectively upregulated in PM or UM indicating that the clinical identities of each patient are reflected in their baseline transcriptional profile (**Figure 2G, Supplementary information 2**). To understand the biological programs associated with these differences, we performed gene ontology analysis on PM-specific and UM-specific upregulated gene sets separately (**Figure 2H**). PM-specific genes were enriched for extracellular matrix organization, blood vessel morphogenesis and negative regulation of both immune responses and WNT signaling, programs collectively consistent with the remodeling of the peritoneal microenvironment and the suppression of anti-tumor immune activity required to establish stable metastatic implants. UM-specific genes, in contrast, were preferentially associated with neuron fate commitment, columnar/cuboidal epithelial differentiation, epithelial cell differentiation and regulation of protein binding, a combination of programs that suggests substantial cell identity plasticity and altered intercellular communication in the chemotherapy-resistant context. Shared enriched terms including animal organ morphogenesis and cell-cell adhesion reflect a common baseline of epithelial organizational capacity retained across all metastatic lines regardless of chemotherapy.

**Figure 2.**
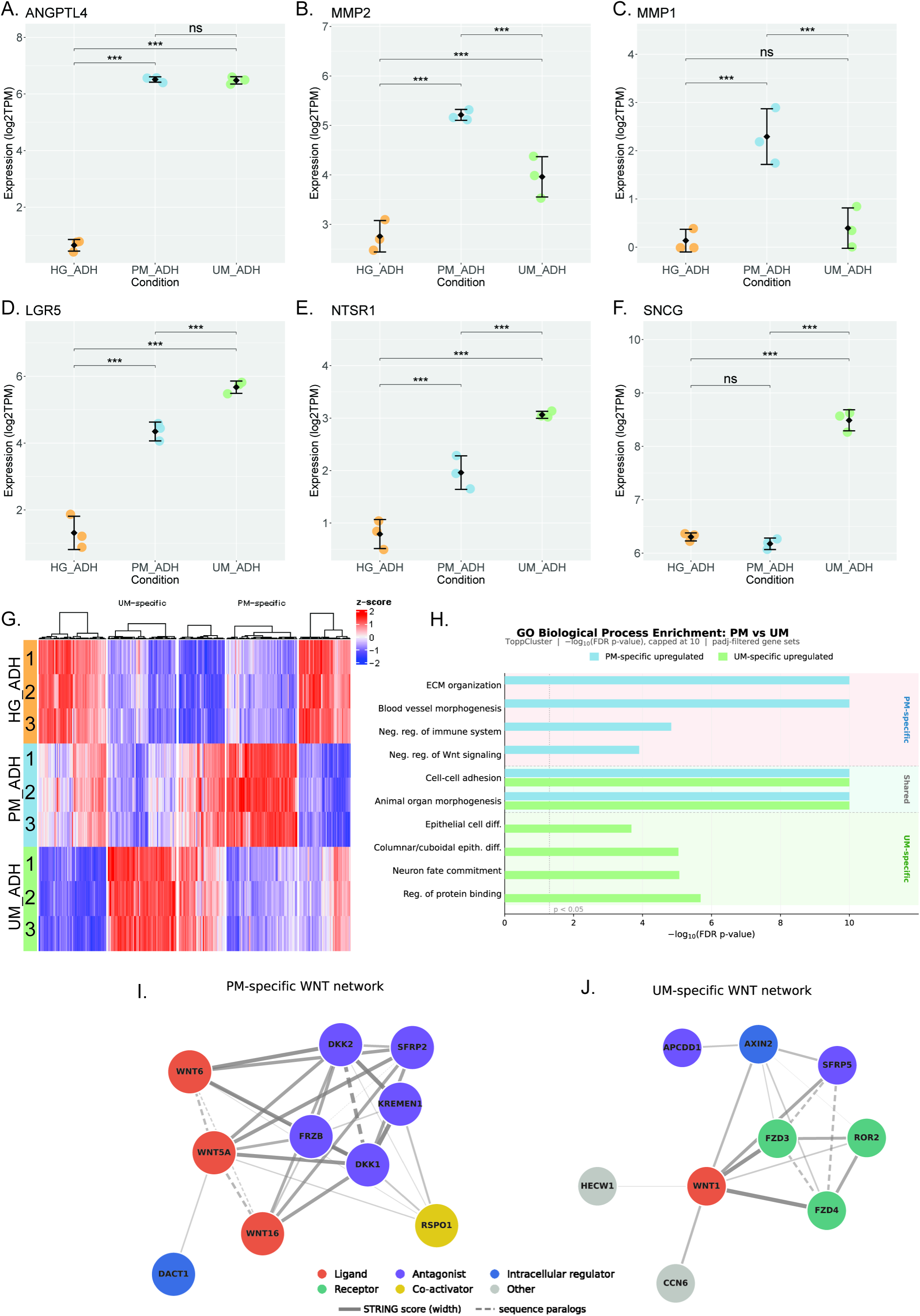
Adherent PDC transcriptomic landscapes. A-F) Expression levels (log₂TPM) of (A-D) metastasisassociated genes (*ANGPTL4, MMP2, MMP1, LGR5*) and (E-F) chemoresistance-associated genes (*NTSR1, SNCG*) across HG, PM, and UM PDCs under adherent culture conditions. Statistical significance was assessed by pairwise comparison; ***p<0.001, ns = not significant. (G) Heatmap (z-score) of genes differentially expressed between PM and HG or between UM and HG in adherent culture, shown across all nine adherent samples (HG, PM, UM; three replicates each). Rows were partitioned by k-means clustering and ordered within each cluster by Ward’s linkage. The PM-specific and UM-specific clusters used for the downstream enrichment and network analyses are indicated. (H) Gene ontology enrichment analysis (GO Biological Process) of PM-specific (blue) and UM-specific (green) upregulated gene sets relative to HG. Dashed line indicates p<0.05 threshold.

To further define the signaling architecture underlying these subtype-specific programs, we applied protein–protein interaction network analysis by using STRING to identify pathway-level hubs within the PM- and UM-specific gene clusters (**Figure 2I-J**). Both networks independently converged on the WNT signaling pathway identifying it as a shared pathway-level feature despite the considerable gene ontology differences described above. It is well documented that both canonical WNT/β-catenin and non-canonical WNT branches have been implicated in ovarian cancer stemness, peritoneal dissemination and acquired chemotherapy resistance [21,28]. Notably, the WNT associated genes identified in PM and UM cells did not overlap and reflected distinct modes of pathway engagement. In PM cells, the upregulated WNT network was dominated by multiple ligands (*WNT5A*, *WNT6* and *WNT16*), the co-activator *RSPO1* and a set of secreted antagonists including *DKK1*, *DKK2*, *SFRP2*, *KREMEN1* and *FRZB*, suggesting a complex autocrine/paracrine WNT environment in which both activating and restraining signals are simultaneously induced (**Figure 2I**). In contrast, the UM-specific WNT network featured the receptors *FZD3*, *FZD4* and *ROR2*, engaged by a single ligand, *WNT1*, and restrained by the secreted antagonist *SFRP5* and the membrane-associated inhibitor *APCDD1*. Because *AXIN2* and *APCDD1* are both direct transcriptional targets of canonical WNT/β-catenin signaling and act as negative feedback regulators, their induction indicates that the canonical branch is actively engaged in UM cells, while the presence of ROR2 indicates a concurrent non-canonical WNT/ROR2 arm associated with invasive and drug-resistant states (**Figure 2J**). This distinction establishes the first of several recurring instances in this study where the three OV-PDCs arrive at the same pathway destination through entirely distinct transcriptional implementations.

### Gene Ontology Pathway Enrichment Analysis defines 3 shared outcomes across all CSC subtypes

Having confirmed that the three OV-PDCs carry distinct baseline transcriptional programs, we next examined how CSC induction reshapes each lineage. To do this, we performed differential expression analysis between SPH and ADH within each OV-PDC, which revealed extensive, line-specific transcriptional changes rather than a shared, uniform CSC signature (**Supplementary Figure 2A-C, Supplementary information 3**). Intersection of CSC-specific upregulated gene sets across HG, PM and UM revealed 34 genes consistently induced in all three SPH cultures relative to matched ADH controls. Meanwhile, hundreds of additional upregulated genes in SPH compared to ADH were unique to individual OV-PDCs (**Figure 3A**). First, the signatures and biological implications of the 34 overlapping genes were evaluated. GO Pathway Enrichment Analysis of the molecular functions, biological processes and cellular components significantly associated within this gene set revealed three groups of interest (**Figure 3B-D**) [29,30]. The molecular function and cellular component enrichment dot plots reveal significant association with components of the extracellular matrix and growth factor binding and signaling (**Figure 3B-C**). Meanwhile, the biological processes enrichment dot plot reveals significant involvement with components of the immune system (**Figure 3D**). Major components of the ECM include collagen and glycosaminoglycans (GAGs), which can exist as components of proteoglycans or independently as hyaluronan [31]. Molecular function enrichment of the 34 shared genes showed a strong association with GAG binding. Since GAGs are negatively charged, they can bind tightly to proteins - including growth factors and cytokines - and influence their availability within the tumor microenvironment [32]. Further, cellular component enrichment showed multiple collagen-associated terms. Collagen is a major structural component of the ECM that provides tissue architecture and support. Cancer cells can utilize collagen for processes such as cell adhesion and migration [31]. As a result, the enrichment of GAG binding and collagen-associated processes suggests that these 34 genes may contribute to ECM modeling establishing the tumor microenvironment (TME).

**Figure 3.**
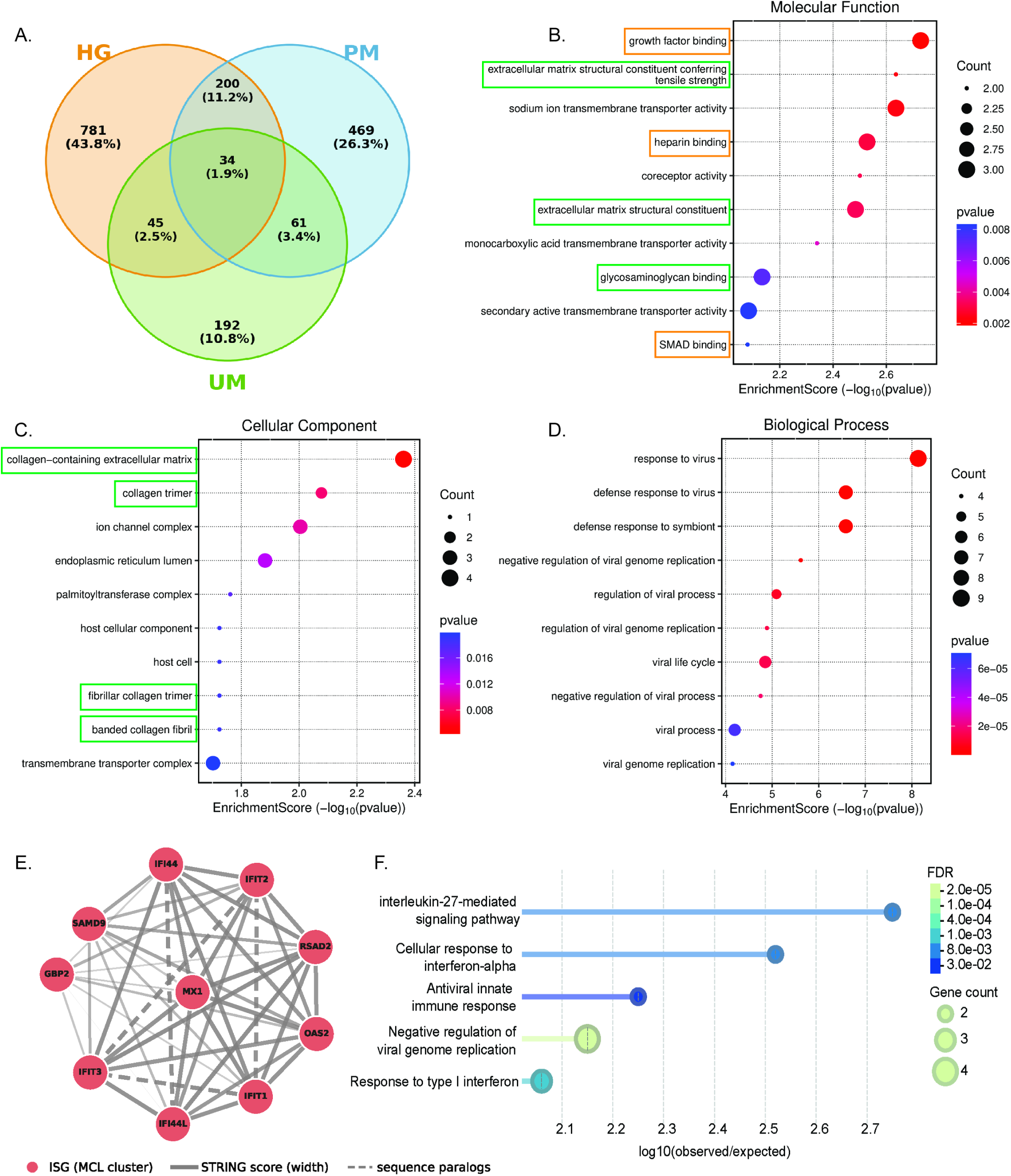
Identification of a conserved cancer stem cell transcriptomic signature across HGSOC PDCs. A) Three-way Venn diagram depicting the overlap of sphere-upregulated genes across HG, PM, and UM PDCs. B-D) GO Pathway Enrichment Analysis of the significant molecular functions (B), cellular components (C), and biological processes (D) of the 34 overlapping genes. Analysis and graphs generated using SRPlot. Gene count represented by dot size and p-value represented by color. Orange boxes represent processes related to growth factor signaling and green boxes represent functions related to the extracellular matrix. E) STRING protein-protein interaction subnetwork of the 34 sph-upregulated genes shared across all 3 PDCs highlighting a densely connected type I interferon/innate immune signaling network. F) Functional enrichment of the interferon-stimulated gene module shown in (E), computed in STRING against a whole-genome background. Gene Ontology Biological Process terms are ordered by false discovery rate (Benjamini– Hochberg-corrected hypergeometric test). Dot size indicates the number of module genes annotated to each term and dot color indicates the FDR; the x axis shows enrichment strength, log_10_ (observed/expected).

### A convergent interferon-stimulated gene module defines a shared transcriptional core across all CSC subtypes

Further analysis of the 34 conserved genes using STRING functional analysis (stringdb.org) revealed one large densely connected network while the rest of the genes either clustered in pairs of 2 or 3 or did not cluster at all (**Supplementary Figure 2D**). This dense network is dominated by immune-related interferon genes. Since the enrichment plot and STRING analysis point to a conserved role of immune-related genes in the 34 overlapping genes, we performed further analysis on their interactions and implications in CSC signaling. Protein– protein interaction analysis showed that these 34 genes form a densely connected network dominated by classical interferon-stimulated genes (ISGs); *MX1*, *IFIT1*/*2*/*3*, *IFI44*/*IFI44L*, *OAS2*, *RSAD2*, *GBP2* and *SAMD9* (**Figure 3E**). Functional enrichment confirmed strong association with “cellular response to interferon-alpha”, “response to type I interferon”, “antiviral innate immune response” and “negative regulation of viral genome replication” (**Figure 3F**). Because these PDC experiments were conducted in the absence of immune or stromal components, this ISG signature cannot be attributed to paracrine cytokine signaling from the tumor microenvironment. Therefore, it most likely reflects a cell-intrinsic, tumor-driven interferon program. So, we propose that the convergent induction of this type I IFN/ISG module reflects a shared adaptive response to cellular stress accompanying CSC formation in ovarian cancer.

### CSC induction reveals patient-specific transcriptional reprogramming

To interpret these changes in the context of established CSC biology, we organized the SPH-upregulated genes into three functional categories that have been repeatedly linked to ovarian CSC behaviors including chemoresistance, metastasis and oncofetal reprogramming and overlaid the HG-CSC, PM-CSC and UM-CSC gene sets onto this framework (**Figure 4A-C**). All three CSC populations activated genes within each of the three functional programs, yet the gene memberships were largely non-overlapping across subtypes indicating that the same hallmarks of CSC identity are built from entirely different molecular components depending on the cell of origin (**Figure 4A-C**). Representative subtype-enriched markers from each category were validated by RT-qPCR, confirming that the RNA-seq-defined patterns are reproducible at the individual gene level (**Supplementary Figure 2E-H**).

**Figure 4.**
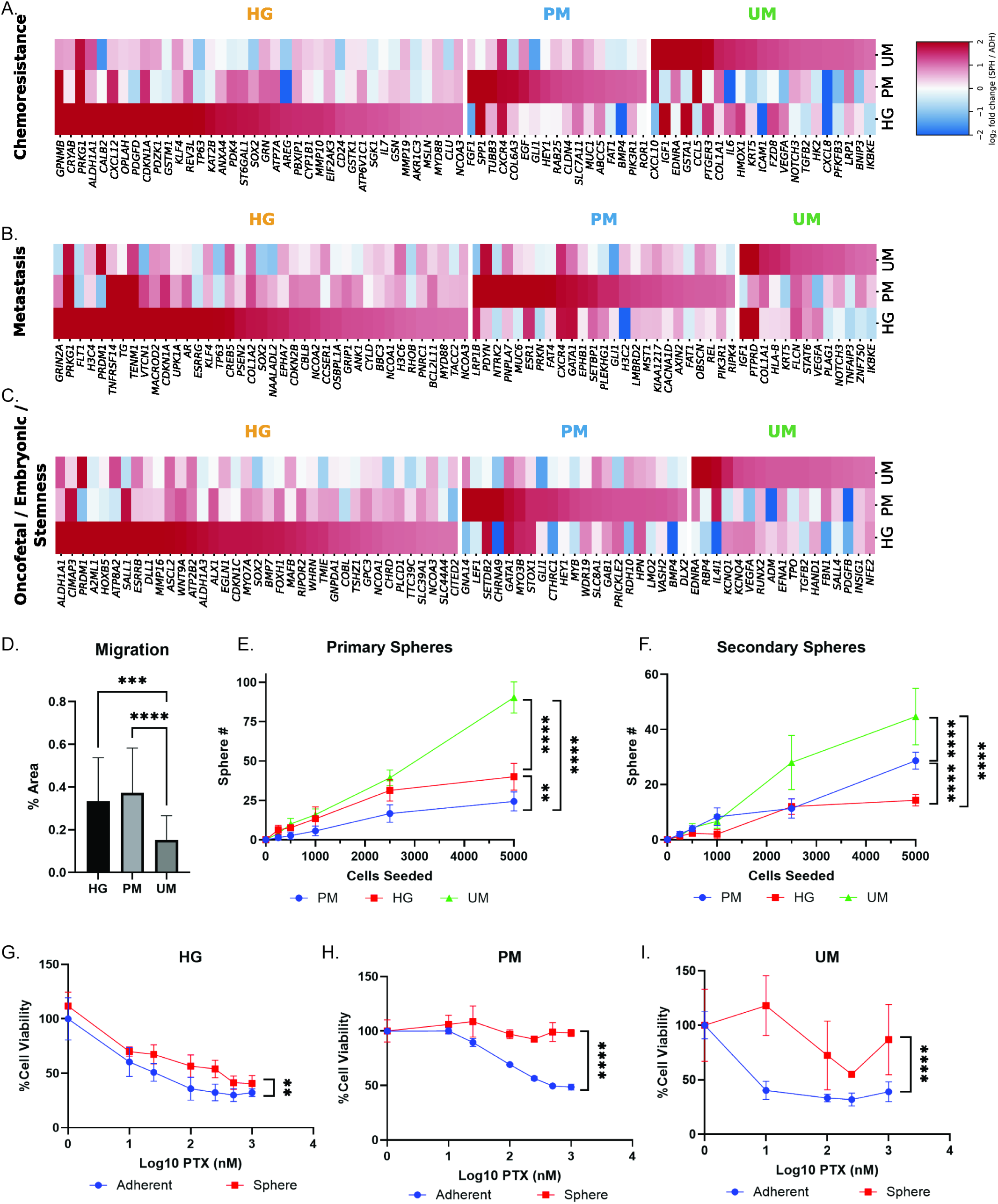
Characterization of distinct cancer stem cell transcriptomic signatures across HGSOC PDCs. A-C) Heatmap of log₂(Sph/Adh) fold change for stemness- and cancer stem cell-associated pathways selected from each PDC Sph-specific upregulated gene set. Genes are organized by expression pattern across HG, PM, and UM. The upper cluster indicates HG-specific genes, middle cluster indicates PM-specificgenes, lower cluster indicates UM-specific genes. Chemoresistance signatures (A), Metastatic signatures (B), and oncofetal/embryonic signatures(C) generated from reference gene sets as described in the methods section. E) Transwell migration assay of SPH from HG, PM and UM. % area of well migrated quantified using imageJ. N=3 experimental and 4 biological replicates. F) Primary sphere formation assay as measure of stemness in HG, PM, UM. N=3 experimental and biological replicates. G) Secondary sphere formation assay as measure of self-renewal in HG, PM, UM. N=3 experimental and biological replicates. H-J) Paclitaxel response assay comparing ADH and SPHs in HG, PM, and UM. Promega 3D Cell Titer Glo measured cell viability and is graphed as a percentage of control viability. Changes in IC50 determined using non-linear regression analyses on GraphPad Prism. N=3 experimental and 4 biological replicates. **p<0.01, ***p<0.001,****p<0.0001.

To validate our findings that while patients’ CSC transcriptomic landscapes are diverse, they share conserved phenotypic implications, we performed a series of functional CSC assays on the three OV-PDCs. First, transwell migrations were performed. All 3 OV-PDC SPHs were capable of migrating through the pores to begin establishing adherent colonies on the basolateral side of the insert (**Figure 4D; Supplementary Figure 3A**). Further, sphere formation assays were performed to evaluate stemness in our three patients. All three were capable of forming spheres reflecting the stemness potential of the PDCs (**Figure 4E**). Next, secondary sphere formation assays were performed to evaluate self-renewal potential of the CSCs, a hallmark feature of stemness. All three PDCs had self-renewal capacity at different rates, and notably PM and UM, the two metastatic PDCs had the highest rates of self-renewal (**Figure 4F; Supplementary Figure 3B**). The migration, sphere formation and self-renewal assays reflect the metastatic and oncofetal/embryonic signatures we see in the transcriptomic data. Finally, since CSCs are intrinsically chemo-resistant, we performed paclitaxel dose response curves comparing ADH and SPH populations in each of the PDCs (**Figure 4G-I; Supplementary Figure 3C**). All three CSCs had significantly higher resistance to paclitaxel than their ADH counterparts. Additionally, PM and UM had markedly higher IC50 shifts compared to HG, likely because UM and PM are metastatic and therefore theoretically more aggressive, and UM is multi-drug resistant. Altogether, the functional assays confirm the results of the transcriptomic signatures. Even though the CSCs between patients have diverse and unique landscapes, their functional outcomes remain the same: stemness, metastasis and resistance.

Next, we evaluated whether these gene-level differences translated into distinct higher-order biological processes. Based on the DEG analysis between CSC and adherent condition on each cell lines, we were able to identify cell type-specific upregulated gene sets (**Supplementary Figure 3D**). Gene ontology analysis of CSC-specific upregulated genes then revealed that HG-CSC is enriched for processes related to neurodevelopment and tissue morphogenesis, PM-CSC for ion channel activity and membrane transport, and UM-CSC for immune and inflammatory responses together with metabolic pathways (**Supplementary Figure 3D**). These complementary lines of evidence indicate that CSC induction doesn’t drive the three OV-PDCs toward a single, convergent molecular state. Instead, each line preserves a memory of its origin and clinical history in the way it assembles stemness, resistance and oncofetal programs. This view is consistent with emerging work showing that ovarian CSC heterogeneity reflects both the epigenetic history of the cell of origin and subsequent therapy-induced reprogramming [22,33]. Collectively, these data support a model in which the canonical hallmarks of cancer stemness including self-renewal, metastatic competence, therapy resistance and acquisition of oncofetal-like traits are presented through distinct transcriptional routes in each OV-PDC subtype. These findings have important implications for therapeutic design. Given that CSC functional properties are realized through distinct gene sets depending on tumor origin, therapeutic strategies targeting a single marker or signaling node may be insufficient to eliminate the full diversity of CSC states encountered across ovarian cancer subtypes.

### Isoform-level reprogramming reveals a hidden layer of subtype-specific CSC identity

Gene-level analysis captures only part of the transcriptional changes that accompany cell-state transitions. To uncover regulatory events that would be missed by gene-centric approaches, we analyzed transcript usage between adherent and CSC conditions in each OV-PDC and systematically searched for significant isoform switches. PM-CSC displayed the highest number of isoform switching events, clearly exceeding those observed in HG-CSC and UM-CSC, indicating that CSC formation in the peritoneal metastasis context is accompanied by particularly extensive remodeling of isoform usage (**Figure 5A**). Importantly, the genes undergoing isoform switching were almost completely distinct among the three lines (**Figure 5B**), extending the subtype specificity seen at the gene level to the isoform layer and demonstrating that CSC reprogramming is stratified across multiple tiers of RNA regulation.

**Figure 5.**
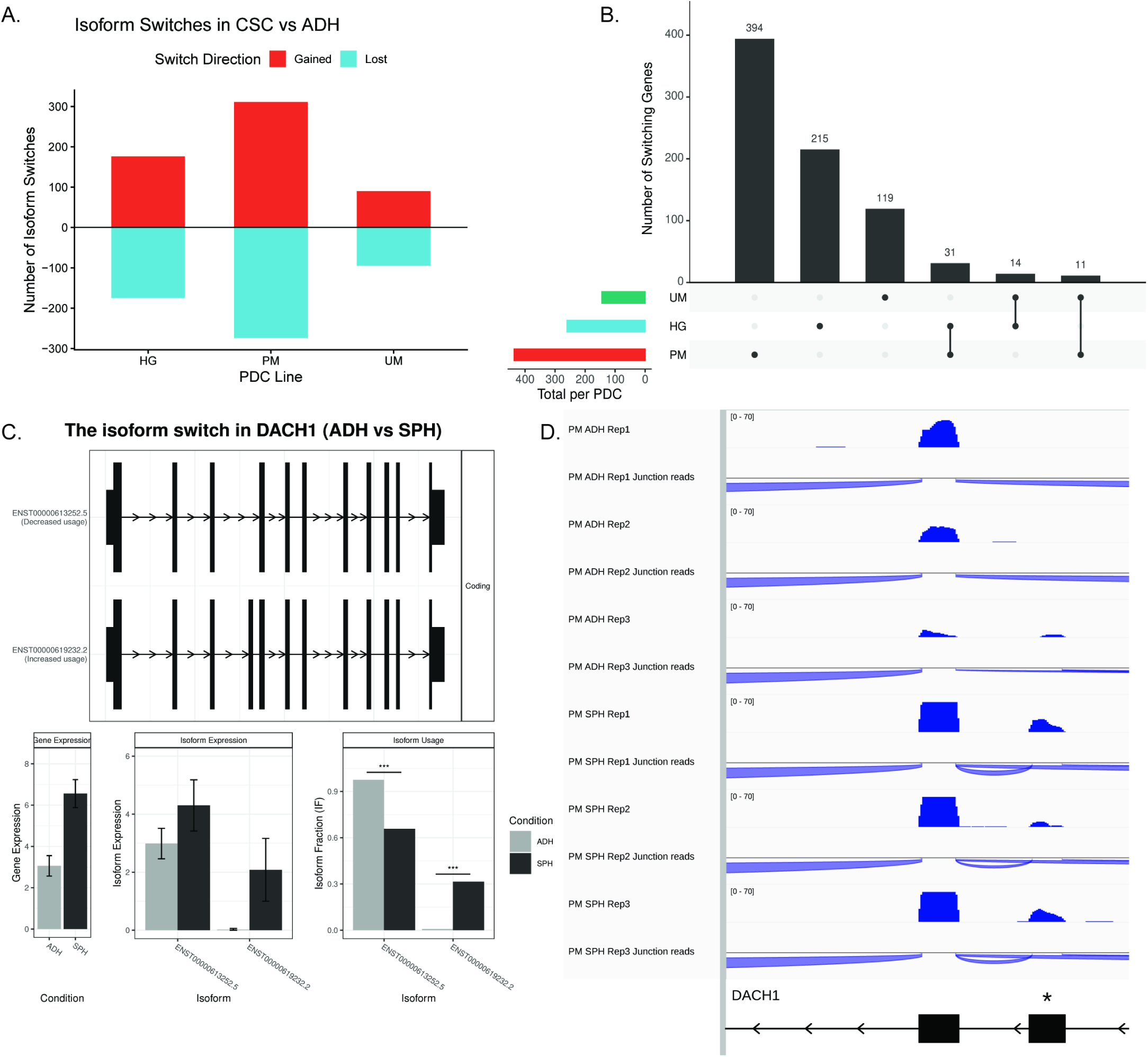
Isoform-level reprogramming reveals subtype-specific CSC identity. A) Number of significant isoform switches between SPH and ADH in each OV-PDC line, separated by direction. Bars above the axis indicate isoforms gained in CSCs and bars below indicate isoforms lost (HG, 176 gained / 175 lost; PM, 311 / 274; UM, 90 / 95). B) UpSet plot of genes undergoing isoform switching. Horizontal bars show the total number of switching genes per PDC and vertical bars show the size of each intersection. C) Isoform switch in *DACH1* between PM-ADH and PM-SPH. Top, exon structures of the two switching isoforms; ENST00000613252.5 decreases in fractional abundance in SPH while ENST00000619232.2 increases, and both are protein coding. Bottom, total *DACH1* gene expression, expression of each isoform, and isoform fraction (IF) in ADH versus SPH. ***q < 0.001. D) Genome browser tracks of *DACH1* in PM-ADH and PM-SPH replicates showing read coverage and splice junction reads, all on the same scale (0–70). The asterisk marks the exon that is differentially used between the two isoforms.

*DACH1* provided a mechanistically informative example of this phenomenon. *DACH1* is a cell-fate determination factor with well-established tumor suppressor functions: it represses the EMT master regulator SNAI1, attenuates YB-1-mediated oncogenic transcription and translation, and its loss of expression has been associated with invasion and poor metastasis-free survival in several cancer types [34,35]. In PM-CSC, total *DACH1* transcript abundance increased relative to PM-ADH, but the ADH-dominant isoform ENST00000613252.5 lost fractional abundance while the alternative isoform ENST00000619232.2 expanded to account for roughly one-third of all *DACH1* transcripts (**Figure 5C**). This pattern was further confirmed by visual inspection of exon usage in genome browser tracks (**Figure 5D**). Thus, *DACH1* is not transcriptionally silenced as a tumor suppressor during CSC induction. Instead, its transcript pool is redistributed between structurally distinct isoforms. Structural annotation showed that both *DACH1* isoforms retain an identical SKI/DAC domain (aa 173–281) indicating that the DNA-binding core and canonical Dach family motif are preserved in both transcripts. The CSC-associated ENST00000619232.2 isoform, however, contains an additional 52-amino acid exon positioned immediately downstream of the SKI/DAC domain within an intrinsically disordered region and shifts the C-terminal coiled-coil segments by 52 residues.

Intrinsically disordered and coiled-coil regions are key hubs for protein-protein interactions and post-translational regulation. Based on these findings, we suggest that the emergence of an isoform with an extended disordered segment and repositioned coiled-coil domains in PM-CSC influence the interaction landscape and regulatory wiring of DACH1. Taken together, these observations describe how isoform-level reprogramming adds a previously hidden layer of subtype-specific CSC identity. Each OV-PDC not only deploys distinct sets of genes to build its CSC state but also reconfigures the isoform composition of key regulatory genes in a subtype-restricted manner.

## 4. Discussion

This study reveals substantial patient-to-patient heterogeneity in the molecular programs that support ovarian CSC states. Using three clinically distinct patient-derived HGSOC models, we found that CSC enrichment reproducibly produced stem-like properties, including sphere-forming and self-renewal, migratory capacity, and increased resistance to paclitaxel compared to adherent cells. However, these shared phenotypic properties were supported by markedly different transcriptional programs. From a clinical perspective, this heterogeneity is particularly relevant to HGSOC, where patients with the same histologic diagnosis can differ substantially in disease progression, treatment response, and development of drug resistance [5,36,37]. Our findings suggest that these clinically important phenotypes may arise through distinct molecular routes rather than a uniform CSC program.

We identified that the shared transcriptional landscape across the three OV-PDCs correlated strongly with immune signaling, ECM organization and growth factor activity. Of the 34 genes that were shared between all three patients and significantly upregulated in CSCs, the largest gene network was the type I interferon/interferon-stimulated-gene signature. This pattern suggests that CSCs may engage in IFN-related programs that influence tumor-cell state and treatment response. Prior studies have linked suboptimal type I IFN signaling to a stem-like and therapy-resistant state through the histone demethylase KDM1B [38,39]. Other studies have shown that ISG-high tumor cell states associated with chronic interferon exposure can confer resistance to both conventional chemotherapy and immune checkpoint blockade through STAT1-dependent epigenomic remodeling [40–42].

The extracellular matrix (ECM) is a supramolecular network of macromolecules, proteins, and glycoproteins that provides scaffolding and organization for cells that are essential for cellular function [6,31,43]. Structurally, the ECM provides a 3D meshwork for tissue architecture, allowing cell adhesion and organization to occur, and is composed of various collagen types, fibronectin, laminin, glycoproteins, proteoglycans, and other structural proteins [6,31,43]. Aside from its structural role, the ECM also plays a large role in essential cell signaling, acting as an intermediary through receptors such as integrins or discoidin domain receptors [31]. These signals help maintain healthy cell processes such as proliferation, differentiation, and overall cell survival [31]. Cancer cells regularly exploit the ECM to support tumor progression. For example, cancer cells utilize the ECM’s structural components to support tumor cell adhesion, migration, and invasion [44]. ECM remodeling can also alter intracellular signaling pathways. For example, integrins, which mediate signaling between the ECM and cells, can sense changes caused by ECM remodeling, including increased matrix stiffness, a common feature of cancer, and activate intracellular signaling pathways that aid in cancer cell proliferation and survival [45]. Beyond supporting general tumor progression, ECM remodeling can also cultivate an environment that supports CSCs [46]. Our data suggest that genes involved in collagen remodeling and ECM structural strength are increased. These changes may contribute to ECM rearrangement and increased matrix stiffness through mechanisms such as collagen crosslinking, ultimately creating a stiffer CSC niche [45]. Increased matrix stiffness can, in turn, promote cancer cell stemness [45]. In addition to promoting cancer stemness, increased ECM stiffness and collagen remodeling can contribute to therapeutic resistance through propagation of chemo resistant cells and limiting effective drug delivery within ovarian tumors [47–49]. The effects of this remodeling extend beyond CSC stemness, as changes in the ECM can also limit immune-cell infiltration [43]. In addition to the structural and mechanical advantages provided by ECM remodeling, the ECM can regulate growth factor availability within the TME, providing another mechanism that may support CSC maintenance.

Growth factors regulate CSCs by increasing stemness properties such as self-renewal, de-differentiation and therapy resistance. As discussed previously, growth factor signaling can interact with inflammatory and ECM cues to promote metastasis and tumor progression. An example of this dynamic interplay is Hepatocyte Growth Factor (HGF) signaling through c-Met in ovarian CSCs. HGF activated by c-met acts in the ascites microenvironment of ovarian cancer to respond to shear stress through regulating the PI3K/AKT pathway thus promoting stemness, metastasis and chemoresistance [50]. Notably, the ECM supports metastatic cell survival under shear stress [51], further underscoring the interconnectivity of this network.

While the conserved gene set provides insight into commonly regulated pathways across ovarian CSCs, the patient-specific transcriptomics provided equally important insight into the landscape of CSCs. HG-CSC upregulated a stemness program centered on epithelial self-renewal and innate immune response simultaneously including the pluripotency factor *SOX2*, the CSC surface marker *CD24* and the NF-κB activator *MYD88.* This is consistent with a stem-like state that is maintained through reinforcement of the original epithelial identity rather than overt plasticity [52]. PM-CSC, in contrast, engaged a developmental morphogen axis driven by Hedgehog effector *GLI1*, the non-canonical WNT receptor *ROR1* and the NOTCH target *HEY1* together with NOTCH/WNT transcriptional mediators *LEF1* and *BMP4*. This configuration collectively programs a high migratory capacity, resistance and EMT-like plasticity suited for peritoneal colonization [53–56]. UM-CSC showed a distinct program in which the oncofetal transcription factors *SALL4* and *RUNX2*, both associated with invasive behavior in ovarian carcinoma, act alongside the inflammatory mediator *CXCL8*, an autocrine driver of platinum and taxane resistance. This combination is consistent with a stress-adapted, inflammation-coupled stemness state in the chemotherapy-exposed line [57,58]. Altogether, the chemoresistance, metastasis and embryonic/oncofetal heatmaps depicting the heterogeneous transcriptomic landscapes of ovarian CSCs reveal how the CSCs maintain phenotypic consistency even with diverse genetic backgrounds.

The implications of our findings span from insight into the conserved signature of CSCs in the ECM, TME, and interferon pathways to the diverse landscape leading to drug resistance, metastasis and upregulation of embryonic pathways. It highlights the importance of delineating both conserved and patient-specific transcriptional signatures to better understand the role of CSCs in ovarian cancer progression. Additionally, we present the importance of deconstructing the heterogeneous signatures between patients. The classic stemness markers do not appear to be conserved across all three patients, illustrating the importance of creative, personalized therapies to combat the phenotypic outcomes of CSCs. While the impact of this study may shift our understanding of CSC signatures, it is not without limitations. This study is limited by sample size and would benefit from future studies with increased patient samples to characterize potential “subtypes” of CSC landscapes between patients and to comprehensively analyze shared and distinct gene signatures between patients. Furthermore, here we have analyzed the shared and diverse upregulated genes amongst samples. An important next step is to evaluate the downregulated signature to identify which genes are turned off in CSCs and what the roles of those genes may be. Future studies evaluating ATAC-sequencing or Hi-C analysis to evaluate epigenetic alterations between CSCs would provide a multiomic picture of the patient derived CSCs. The data presented here should also be further evaluated in functional studies to determine which factors are critical modulators of CSCs verses byproducts of CSC upregulation. Finally, based on the heterogeneous signature from patient to patient, it is possible that the CSC subpopulation itself consists of multiple subpopulations of CSCs which each regulate different phenotypic outcomes in the tumor microenvironment. Therefore, single-cell sequencing approaches would bridge this gap to identify clustering within CSCs. Altogether, this paper presents both conserved and distinct transcriptomic signatures between three patient derived ovarian cancer samples providing insight into CSC signaling, diversity and future applications for personalized therapeutic approaches.

## 5. Conclusions

Through comparing CSCs with ADH samples from 3 different OV-PDCs, we found that CSC induction reorganized the transcriptome across every OV-PDC, yet only 34 upregulated genes were shared by all 3 patients. These genes pointed to extracellular matrix remodeling, growth factor binding, and a type I interferon-stimulated gene network as a conserved network amongst OV-PDC CSCs. In contrast, the chemoresistance, metastasis, and oncofetal programs were all upregulated in CSCs, but were comprised of largely non-overlapping genes in each OV-PDC, while isoform switching was almost entirely patient-specific. Functionally, all 3 OV-PDCs showed comparable migration, self-renewal, and paclitaxel resistance, indicating that conserved CSC phenotypes arise through patient-specific molecular routes. In summary our findings represent both conserved and distinct transcriptomic signatures between CSCs from patient derived HGSOC samples revealing patient heterogeneity within CSC signaling and highlighting the importance for novel personalized therapeutic approaches.

## Supplementary Materials

The following supporting information is available: Supplementary Figure 1: 3D Principal Component Analysis and RT-qPCR Validation; Supplementary Figure 2: Cancer stem cell transcriptomic signature and validation in OV-PDCs; Supplementary Figure 3: Additional data for functional assays and distinct biological processes. Supplementary Table 1: Primer list; Supplementary File 1: PCA loadings; Supplementary File 2: ADH baseline clusters; Supplementary File 3: SPH vs ADH DEGs.

## Author Contributions

Conceptualization: WYC & RSP. Ideas: WYC & RSP. Data curation: MSJ, LJD, AAB, JK, CF, AP, WYC, RSP. Formal analysis: MSJ, LJD, AAB, JK, CF, AP, WYC, RSP. Funding acquisition: WYC & RSP. Investigation: MSJ, LJD, AAB, JK, CF, AP, WYC & RSP. Methodology: SCL, WZ, WYC & RSP. Project administration: WZ, WYC & RSP. Resources: SCL, WZ, WYC & RSP. Software: JK & WYC. Supervision: WZ, WYC & RSP. Validation: MSJ, LJD, AAB & RSP. Visualization: MSJ, LJD, AAB, JK, WYC & RSP. Writing – original draft: MSJ, LJD, AAB, JK, SD, JY, RPM, WZ, WYC & RSP. Writing – review & editing: MSJ, LJD, AAB, SD, JY, SCL, RPM, WZ, WYC & RSP.

## Institutional Review Board Statement

Not applicable

## Informed Consent Statement

Not applicable

## Data Availability Statement

The datasets analyzed in this study are uploaded in the GEO database (GSE348927).

## Acknowledgments

The authors thank the UTHSC Center for Cancer Research Translational Science Shared Resource for assistance in RNA-sequencing of samples.

## Abbreviations

The following abbreviations are used in this manuscript:

ACTB: Beta-Actin
ADH: Adherent
CSCs: Cancer stem cells
DEG: Differentially expressed gene
ECM: Extracellular matrix
EMT: Epithelial-to-mesenchymal transition
GAG: Glycosaminoglycan
HG: Primary HGSOC sample
HGSOC: High grade serous ovarian cancer
IFN: Interferon
ISG: Interferon-stimulated gene
OV-PDC: Ovarian cancer patient derived cells
PDCs: Patient-derived ovarian cancer cells
PD-HGSOC: Patient-derived high grade serous ovarian cancer
PDMR: Patient derived material repository
PM: Peritoneal metastatic HGSOC sample
PTX: Paclitaxel
SPH: 3D-CSC-enriched spheroids
TME: Tumor microenvironment
TPM: Transcript-per-million
UM: Peritoneal metastatic HGSOC from umbilical biopsy sample

## Supplementary Figures

**Supplementary Figure 1.**
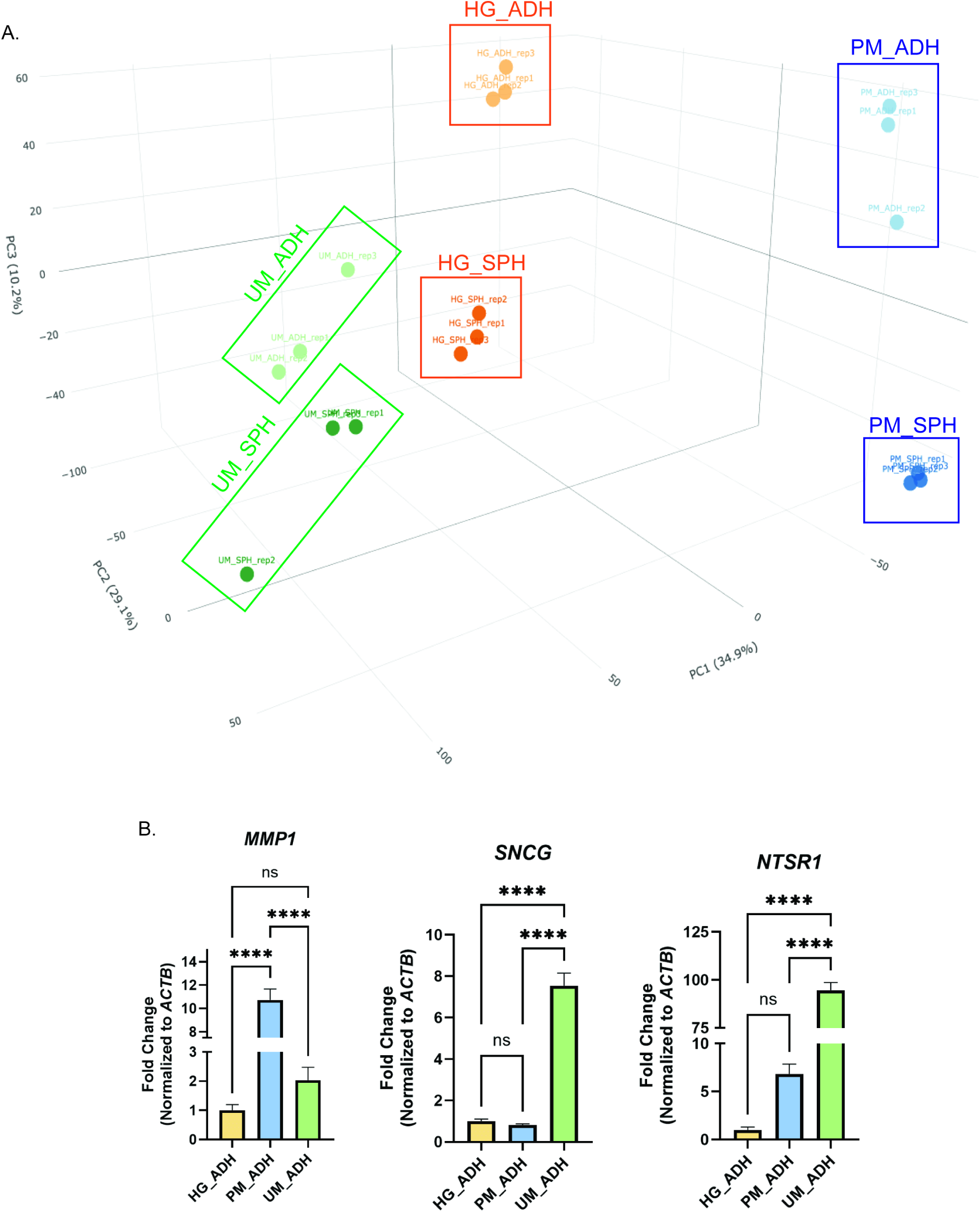
3D Principal Component Analysis and RT-qPCR Validation. A) 3-Dimmensional PCA of RNA-seq data from all 9 groups showing transcriptomic separation between ADH and SPH conditions and between patients in HG, PM, and UM cells. HG = orange, PM = blue, UM = green. B) RT-qPCR validation of *MMP1, SNCG*, and *NTSR1* expression normalized to *ACTB* across HG_ADH, PM_ADH, and UM_ADH conditions, confirming the RNA-seq findings at the transcript level.

**Supplementary Figure 2.**
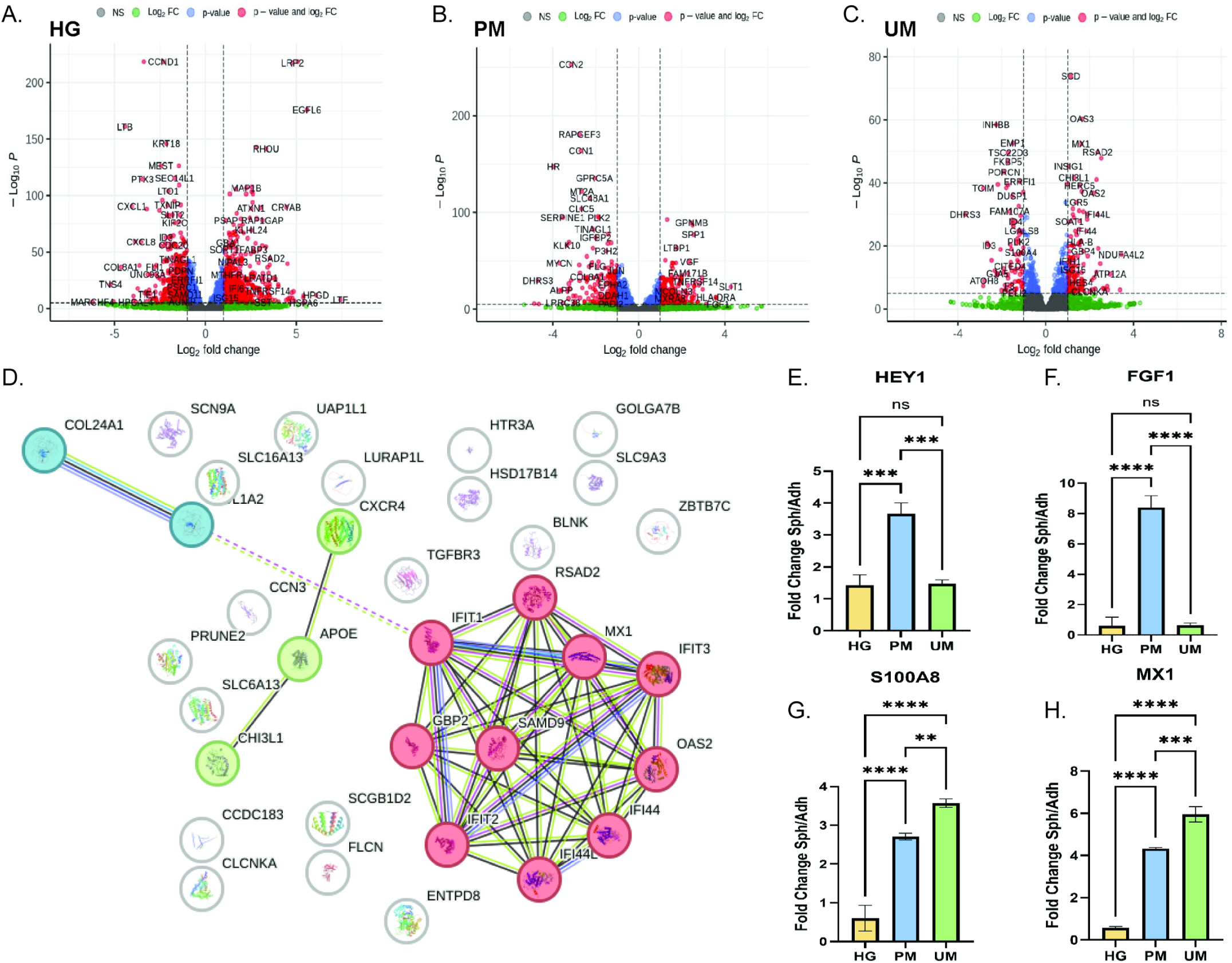
Cancer stem cell transcriptomic signature and validation in OV-PDCs. A-C) Volcano plots differential expression analysis comparing SPH versus ADH conditions in HG(A), PM (B), and UM (C) PDCs. Genes are colored by significance criteria: red = significant by both p-value and log₂ fold change; blue = log₂FC only; green =p-value only; gray = not significant. Dashed lines indicate log₂FC= ±1 and adjusted p-value = 0.05 thresholds. D) STRING Network analysis of all 34 genes highlighting 1 dense cluster, 1 cluster of 3 proteins and 1 cluster of 2 proteins while the rest do not cluster together. Generated using stringdb.org E-H) RT-qPCR validation of representative CSC-associated genes, expressed as fold change (Sph/Adh) normalized to ACTB, across HG, PM, and UM PDCs. (E) HEY1 and (F) FGF1 are preferentially upregulated in PM spheres. (G) S100A8 is most prominently induced in UM spheres. (H) MX1 is upregulated across all three PDC types in sphere conditions.**p<0.01,***p<0.001, ****p<0.0001, ns = not significant.

**Supplementary Figure 3:**
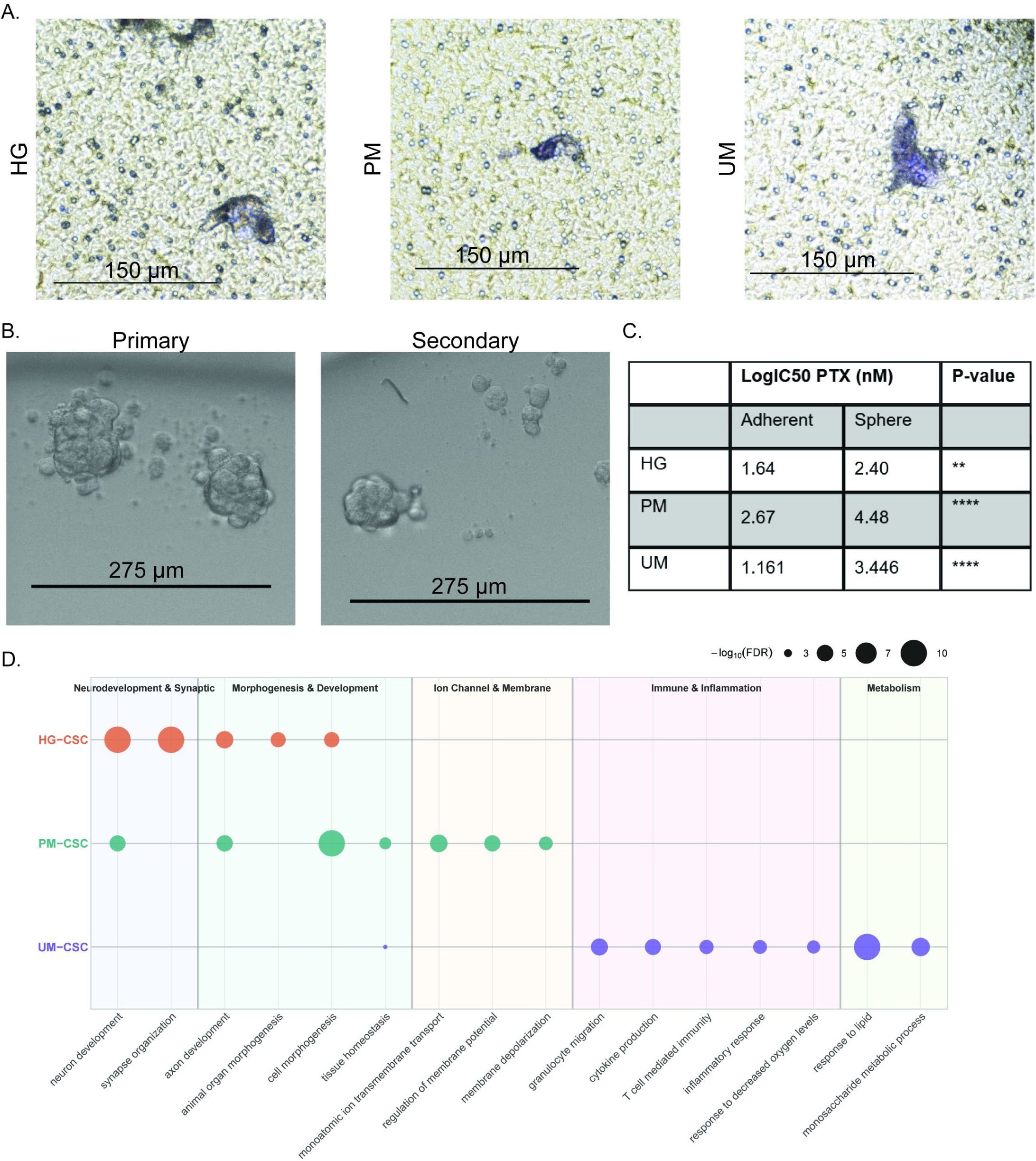
Additional data for functional assays and distinct biological processes in CSCs. A) Representative images of migrated cells from CSCs. Scale bar= 150 µm. B) Representative images of primary vs secondary spheres from sphere formation and self-renewal assays. Scalebar = 275 µm. C) Table of log10 IC50 of paclitaxel (PTX) in ADH vs SPH for HG, PM, and UM. **p<0.01, ****p<0.0001. D) Bubble plot of GO Biological Process enrichment for PDC-specific sphere-upregulated gene sets. Bubble size reflects−log₁₀(FDR).

**Supplementary Table 1.**
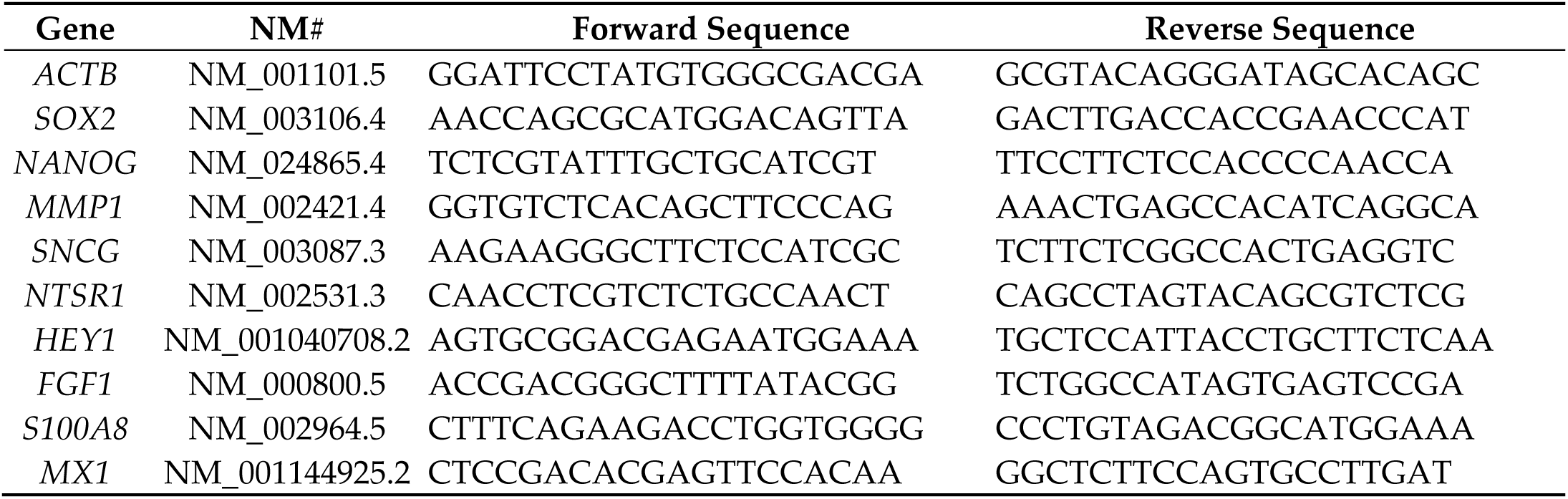
Primer List.

## References

[1] Siegel RL, Kratzer TB, Wagle NS, Sung H, Jemal A. Cancer statistics, 2026. CA Cancer J Clin 2026;76:e70043. 10.3322/caac.70043.

[2] Smolarz B, Biernacka K, Łukasiewicz H, Samulak D, Piekarska E, Romanowicz H, et al. Ovarian Cancer-Epidemiology, Classification, Pathogenesis, Treatment, and Estrogen Receptors’ Molecular Backgrounds. Int J Mol Sci 2025;26. 10.3390/ijms26104611.

[3] Yang L, Xie H-J, Li Y-Y, Wang X, Liu X-X, Mai J. Molecular mechanisms of platinum-based chemotherapy resistance in ovarian cancer (Review). Oncol Rep 2022;47. 10.3892/or.2022.8293.

[4] Richardson DL, Eskander RN, O’Malley DM. Advances in Ovarian Cancer Care and Unmet Treatment Needs for Patients With Platinum Resistance: A Narrative Review. JAMA Oncol 2023;9:851–9. 10.1001/jamaoncol.2023.0197.

[5] Azzalini E, Stanta G, Canzonieri V, Bonin S. Overview of Tumor Heterogeneity in High-Grade Serous Ovarian Cancers. Int J Mol Sci 2023;24. 10.3390/ijms242015077.

[6] S. Perkins R, Choi W-Y, S. Jung and Wenjing Zhang M. Tumor Microenvironment of Epithelial Ovarian Cancer, 2025. 10.5772/intechopen.1013212.

[7] Lee H, Kim B, Park J, Park S, Yoo G, Yum S, et al. Cancer stem cells: landscape, challenges and emerging therapeutic innovations. Signal Transduct Target Ther 2025;10:248. 10.1038/s41392-025-02360-2.

[8] Manni W, Min W. Signaling pathways in the regulation of cancer stem cells and associated targeted therapy. MedComm (Beijing) 2022;3:e176. 10.1002/mco2.176.

[9] Phi LTH, Sari IN, Yang Y-G, Lee S-H, Jun N, Kim KS, et al. Cancer Stem Cells (CSCs) in Drug Resistance and their Therapeutic Implications in Cancer Treatment. Stem Cells Int 2018;2018:5416923. 10.1155/2018/5416923.

[10] Chu X, Tian W, Ning J, Xiao G, Zhou Y, Wang Z, et al. Cancer stem cells: advances in knowledge and implications for cancer therapy. Signal Transduct Target Ther 2024;9:170. 10.1038/s41392-024-01851-y.

[11] Lee J, Kotliarova S, Kotliarov Y, Li A, Su Q, Donin NM, et al. Tumor stem cells derived from glioblastomas cultured in bFGF and EGF more closely mirror the phenotype and genotype of primary tumors than do serum-cultured cell lines. Cancer Cell 2006;9:391–403. 10.1016/j.ccr.2006.03.030.

[12] Bapat SA, Mali AM, Koppikar CB, Kurrey NK. Stem and progenitor-like cells contribute to the aggressive behavior of human epithelial ovarian cancer. Cancer Res 2005;65:3025–9. 10.1158/0008-5472.CAN-04-3931.

[13] Ajith AK, Subramani S, Manickam AH, Ramasamy S. Chemotherapeutic Resistance Genes of Breast Cancer Patients - An Overview. Adv Pharm Bull 2022;12:649–57. 10.34172/apb.2022.048.

[14] Kang HC, Kim I-J, Park J-H, Shin Y, Ku J-L, Jung MS, et al. Identification of genes with differential expression in acquired drug-resistant gastric cancer cells using high-density oligonucleotide microarrays. Clin Cancer Res 2004;10:272–84. 10.1158/1078-0432.ccr-1025-3.

[15] Huang D, Savage SR, Calinawan AP, Lin C, Zhang B, Wang P, et al. A highly annotated database of genes associated with platinum resistance in cancer. Oncogene 2021;40:6395–405. 10.1038/s41388-021-02055-2.

[16] Ramaswamy S, Ross KN, Lander ES, Golub TR. A molecular signature of metastasis in primary solid tumors. Nat Genet 2003;33:49–54. 10.1038/ng1060.

[17] Dymerska D, Marusiak AA. Drivers of cancer metastasis – Arise early and remain present. Biochimica et Biophysica Acta (BBA) - Reviews on Cancer 2024;1879:189060. 10.1016/j.bbcan.2023.189060.

[18] Bruggeman JW, Irie N, Lodder P, van Pelt AMM, Koster J, Hamer G. Tumors Widely Express Hundreds of Embryonic Germline Genes. Cancers (Basel) 2020;12. 10.3390/cancers12123812.

[19] Wong DJ, Liu H, Ridky TW, Cassarino D, Segal E, Chang HY. Module map of stem cell genes guides creation of epithelial cancer stem cells. Cell Stem Cell 2008;2:333–44. 10.1016/j.stem.2008.02.009.

[20] Nguyen A, Lausten M, Boman BM. Oncofetal Reprogramming: A New Frontier in Cancer Therapy Resistance. International Journal of Translational Medicine 2026;6:6. 10.3390/ijtm6010006.

[21] Teeuwssen M, Fodde R. Wnt Signaling in Ovarian Cancer Stemness, EMT, and Therapy Resistance. J Clin Med 2019;8. 10.3390/jcm8101658.

[22] Hatina J, Boesch M, Sopper S, Kripnerova M, Wolf D, Reimer D, et al. Ovarian Cancer Stem Cell Heterogeneity. Adv Exp Med Biol 2019;1139:201–21. 10.1007/978-3-030-14366-4_12.

[23] Li Y-K, Gao A-B, Zeng T, Liu D, Zhang Q-F, Ran X-M, et al. ANGPTL4 accelerates ovarian serous cystadenocarcinoma carcinogenesis and angiogenesis in the tumor microenvironment by activating the JAK2/STAT3 pathway and interacting with ESM1. J Transl Med 2024;22:46. 10.1186/s12967-023-04819-8.

[24] Yokoi A, Yoshioka Y, Yamamoto Y, Ishikawa M, Ikeda S-I, Kato T, et al. Malignant extracellular vesicles carrying MMP1 mRNA facilitate peritoneal dissemination in ovarian cancer. Nat Commun 2017;8:14470. 10.1038/ncomms14470.

[25] Mizutani K, Kofuji K, Shirouzu K. The significance of MMP-1 and MMP-2 in peritoneal disseminated metastasis of gastric cancer. Surg Today 2000;30:614–21. 10.1007/s005950070101.

[26] Liu W, Zhang J, Gan X, Shen F, Yang X, Du N, et al. LGR5 promotes epithelial ovarian cancer proliferation, metastasis, and epithelial-mesenchymal transition through the Notch1 signaling pathway. Cancer Med 2018;7:3132–42. 10.1002/cam4.1485.

[27] Wang W, Lokman NA, Barry SC, Oehler MK, Ricciardelli C. LGR5: An emerging therapeutic target for cancer metastasis and chemotherapy resistance. Cancer Metastasis Rev 2025;44:23. 10.1007/s10555-024-10239-x.

[28] Barbolina M V, Burkhalter RJ, Stack MS. Diverse mechanisms for activation of Wnt signalling in the ovarian tumour microenvironment. Biochem J 2011;437:1–12. 10.1042/BJ20110112.

[29] Luo W, Brouwer C. Pathview: an R/Bioconductor package for pathway-based data integration and visualization. Bioinformatics 2013;29:1830–1. 10.1093/bioinformatics/btt285.

[30] Tang D, Chen M, Huang X, Zhang G, Zeng L, Zhang G, et al. SRplot: A free online platform for data visualization and graphing. PLoS One 2023;18:e0294236. 10.1371/journal.pone.0294236.

[31] Naba A. Mechanisms of assembly and remodelling of the extracellular matrix. Nat Rev Mol Cell Biol 2024;25:865–85. 10.1038/s41580-024-00767-3.

[32] Silva JC, Carvalho MS, Han X, Xia K, Mikael PE, Cabral JMS, et al. Compositional and structural analysis of glycosaminoglycans in cell-derived extracellular matrices. Glycoconj J 2019;36:141–54. 10.1007/s10719-019-09858-2.

[33] Eun K, Ham SW, Kim H. Cancer stem cell heterogeneity: origin and new perspectives on CSC targeting. BMB Rep 2017;50:117–25. 10.5483/bmbrep.2017.50.3.222.

[34] Wu K, Chen K, Wang C, Jiao X, Wang L, Zhou J, et al. Cell fate factor DACH1 represses YB-1-mediated oncogenic transcription and translation. Cancer Res 2014;74:829–39. 10.1158/0008-5472.CAN-13-2466.

[35] Zhao F, Wang M, Li S, Bai X, Bi H, Liu Y, et al. DACH1 inhibits SNAI1-mediated epithelial-mesenchymal transition and represses breast carcinoma metastasis. Oncogenesis 2015;4:e143. 10.1038/oncsis.2015.3.

[36] Veneziani AC, Gonzalez-Ochoa E, Alqaisi H, Madariaga A, Bhat G, Rouzbahman M, et al. Heterogeneity and treatment landscape of ovarian carcinoma. Nat Rev Clin Oncol 2023;20:820–42. 10.1038/s41571-023-00819-1.

[37] Nath A, Cosgrove PA, Mirsafian H, Christie EL, Pflieger L, Copeland B, et al. Evolution of core archetypal phenotypes in progressive high grade serous ovarian cancer. Nat Commun 2021;12:3039. 10.1038/s41467-021-23171-3.

[38] De Martino M, Vanpouille-Box C. Type I interferon induces cancer stem cells-mediated chemotherapy resistance. Oncoimmunology 2022;11:2127274. 10.1080/2162402X.2022.2127274.

[39] Musella M, Guarracino A, Manduca N, Galassi C, Ruggiero E, Potenza A, et al. Type I IFNs promote cancer cell stemness by triggering the epigenetic regulator KDM1B. Nat Immunol 2022;23:1379–92. 10.1038/s41590-022-01290-3.

[40] Qiu J, Xu B, Ye D, Ren D, Wang S, Benci JL, et al. Cancer cells resistant to immune checkpoint blockade acquire interferon-associated epigenetic memory to sustain T cell dysfunction. Nat Cancer 2023;4:43–61. 10.1038/s43018-022-00490-y.

[41] Jacquelot N, Yamazaki T, Roberti MP, Duong CPM, Andrews MC, Verlingue L, et al. Sustained Type I interferon signaling as a mechanism of resistance to PD-1 blockade. Cell Res 2019;29:846–61. 10.1038/s41422-019-0224-x.

[42] Benci JL, Xu B, Qiu Y, Wu TJ, Dada H, Twyman-Saint Victor C, et al. Tumor Interferon Signaling Regulates a Multigenic Resistance Program to Immune Checkpoint Blockade. Cell 2016;167:1540–1554.e12. 10.1016/j.cell.2016.11.022.

[43] Brown Y, Hua S, Tanwar PS. Extracellular matrix in high-grade serous ovarian cancer: Advances in understanding of carcinogenesis and cancer biology. Matrix Biol 2023;118:16–46. 10.1016/j.matbio.2023.02.004.

[44] He X, Lee B, Jiang Y. Extracellular matrix in cancer progression and therapy. Medical Review (2021) 2022;2:125–39. 10.1515/mr-2021-0028.

[45] Safaei S, Sajed R, Shariftabrizi A, Dorafshan S, Saeednejad Zanjani L, Dehghan Manshadi M, et al. Tumor matrix stiffness provides fertile soil for cancer stem cells. Cancer Cell Int 2023;23:143. 10.1186/s12935-023-02992-w.

[46] Brown Y, Hua S, Tanwar PS. Extracellular matrix-mediated regulation of cancer stem cells and chemoresistance. Int J Biochem Cell Biol 2019;109:90–104. 10.1016/j.biocel.2019.02.002.

[47] Sherman-Baust CA, Weeraratna AT, Rangel LBA, Pizer ES, Cho KR, Schwartz DR, et al. Remodeling of the extracellular matrix through overexpression of collagen VI contributes to cisplatin resistance in ovarian cancer cells. Cancer Cell 2003;3:377–86. 10.1016/s1535-6108(03)00058-8.

[48] Zhao Y, Cao J, Melamed A, Worley M, Gockley A, Jones D, et al. Losartan treatment enhances chemotherapy efficacy and reduces ascites in ovarian cancer models by normalizing the tumor stroma. Proc Natl Acad Sci U S A 2019;116:2210–9. 10.1073/pnas.1818357116.

[49] Pietilä EA, Gonzalez-Molina J, Moyano-Galceran L, Jamalzadeh S, Zhang K, Lehtinen L, et al. Coevolution of matrisome and adaptive adhesion dynamics drives ovarian cancer chemoresistance. Nat Commun 2021;12:3904. 10.1038/s41467-021-24009-8.

[50] Hassan AA, Artemenko M, Tang MKS, Shi Z, Chen L-Y, Lai H-C, et al. Ascitic fluid shear stress in concert with hepatocyte growth factor drive stemness and chemoresistance of ovarian cancer cells via the c-Met-PI3K/Akt-miR-199a-3p signaling pathway. Cell Death Dis 2022;13:537. 10.1038/s41419-022-04976-6.

[51] Jalali S, del Pozo MA, Chen K, Miao H, Li Y, Schwartz MA, et al. Integrin-mediated mechanotransduction requires its dynamic interaction with specific extracellular matrix (ECM) ligands. Proc Natl Acad Sci U S A 2001;98:1042–6. 10.1073/pnas.98.3.1042.

[52] Lee SH, Oh S-Y, Do SI, Lee HJ, Kang HJ, Rho YS, et al. SOX2 regulates self-renewal and tumorigenicity of stem-like cells of head and neck squamous cell carcinoma. Br J Cancer 2014;111:2122–30. 10.1038/bjc.2014.528.

[53] Ke Z, Caiping S, Qing Z, Xiaojing W. Sonic hedgehog-Gli1 signals promote epithelial-mesenchymal transition in ovarian cancer by mediating PI3K/AKT pathway. Med Oncol 2015;32:368. 10.1007/s12032-014-0368-y.

[54] Zhang S, Cui B, Lai H, Liu G, Ghia EM, Widhopf GF, et al. Ovarian cancer stem cells express ROR1, which can be targeted for anti-cancer-stem-cell therapy. Proc Natl Acad Sci U S A 2014;111:17266–71. 10.1073/pnas.1419599111.

[55] McAuliffe SM, Morgan SL, Wyant GA, Tran LT, Muto KW, Chen YS, et al. Targeting Notch, a key pathway for ovarian cancer stem cells, sensitizes tumors to platinum therapy. Proc Natl Acad Sci U S A 2012;109:E2939–48. 10.1073/pnas.1206400109.

[56] Thériault BL, Shepherd TG, Mujoomdar ML, Nachtigal MW. BMP4 induces EMT and Rho GTPase activation in human ovarian cancer cells. Carcinogenesis 2007;28:1153–62. 10.1093/carcin/bgm015.

[57] Yang M, Xie X, Ding Y. SALL4 is a marker of poor prognosis in serous ovarian carcinoma promoting invasion and metastasis. Oncol Rep 2016;35:1796–806. 10.3892/or.2016.4545.

[58] Wang Y, Qu Y, Niu XL, Sun WJ, Zhang XL, Li LZ. Autocrine production of interleukin-8 confers cisplatin and paclitaxel resistance in ovarian cancer cells. Cytokine 2011;56:365–75. 10.1016/j.cyto.2011.06.005.

